# The dynamic basis of cognition: an integrative core under the control of the ascending neuromodulatory system

**DOI:** 10.1101/266635

**Authors:** J.M. Shine, M. Breakspear, P.T. Bell, K. Ehgoetz Martens, R. Shine, O. Koyejo, O. Sporns, R.A. Poldrack

## Abstract

The human brain integrates diverse cognitive processes into a coherent whole, shifting fluidly as a function of changing environmental demands. Despite recent progress, the neurobiological mechanisms responsible for this dynamic system-level integration remain poorly understood. Here, we used multi-task fMRI data from the Human Connectome Project to examine the spatiotemporal architecture of cognition in the human brain. By investigating the spatial, dynamic and molecular signatures of system-wide neural activity across a range of cognitive tasks, we show that large-scale neuronal activity converges onto a low dimensional manifold that facilitates the dynamic execution of diverse task states. Flow within this attractor space is associated with dissociable cognitive functions, and with unique patterns of network-level topology and information processing complexity. The axes of the low-dimensional neurocognitive architecture align with regional differences in the density of neuromodulatory receptors, which in turn relate to distinct signatures of network controllability estimated from the structural connectome. These results advance our understanding of functional brain organization by emphasizing the interface between low dimensional neural activity, network topology, neuromodulatory systems and cognitive function.

**One Sentence Summary:** A diverse set of neuromodulators facilitates the formation of a dynamic, low-dimensional integrative core in the brain that is recruited by diverse cognitive demands

## Main Text

The human brain is a complex adaptive system in which diverse behaviors arise from coordinated neural activity across a range of spatial and temporal scales. Linking activity within this large-scale neural architecture to the diversity of cognitive states remains an important goal for neuroscience.

The analysis of complex brain networks can be used to discern the organizational properties of the brain that are crucial for its functional dynamics^1^. Network accounts of brain function have hypothesized that a ‘dynamic core’ of regions flexibly guide the flow of activity within the brain to facilitate cognition^2,3^. These frameworks predict that a distributed set of core regions is active across multiple tasks^4,5^ and integrates more specialized regions^6^, altering baseline communication dynamics in service of task-specific computations. Although computational approaches have investigated these large-scale patterns^8^, little is currently known about the mechanisms that facilitate system-wide brain state dynamics as a function of cognition.

One tractable approach to this problem is to exploit the redundancy within complex systems by utilizing dimensionality reduction techniques^9,10^. These approaches uncover latent functional patterns in complex datasets by distilling brain activity patterns into spatiotemporally similar components^10^. The dynamics of the system can then be interrogated within this low-dimensional space, offering insights into the mechanisms that underlie the system’s function. These approaches have successfully been used in the past to elucidate the functional brain circuitry that underlies the behavioral repertoires of a diverse range of organisms, including the nematode (*Caenorhabditis elegans*), the fruit fly (*Drosophila melanogaster*) and the ferret (*Mustela putorius furo*)^11-14^. Computational modeling suggests that similar dynamic low-dimensional mechanisms should exist within human brains^1^.

Here, we analyze whole-brain functional neuroimaging data across a suite of cognitive tasks to identify the low-dimensional dynamic core of cognition in the human brain. We demonstrate that the dynamic functional organization of the brain across a suite of cognitive tasks describes a flow along a low-dimensional phase space. This dynamic flow aligns with unique cognitive brain states that recur across distinct cognitive tasks. We next demonstrate that the flow of activity reflects an ‘integrative core’ that maximizes information processing complexity over relatively long time scales. Finally, we show that the axes of this low-dimensional space are closely related to spatial patterns of gene expression for specific families of neuromodulatory receptors, and also to unique signatures of structural network controllability, which together provide a plausible biological mechanism for the modulation of system-level brain dynamics. These results present a novel view of brain function based on the coordinated dynamics of functional brain networks over time and provide a mechanistic account of the association between a suite of distinct neuromodulatory systems and cognitive function.

### Low dimensional global brain activity recurs across multiple cognitive tasks

We used high temporal resolution 3T fMRI data (TR = 0.72s) from the Human Connectome Project to examine Blood Oxygen Level Dependent (BOLD) activity from 200 unrelated individuals across seven cognitive tasks, each of which engages distinct cognitive functions: emotional processing, gambling, mathematical calculation, language processing, motor execution, working memory performance, relational matching and social inference^15^. Preprocessed BOLD time series were extracted from 375 cortical and subcortical parcels^16^ and concatenated across all seven tasks and across all subjects. To ensure reproducibility, analyses were initially developed within an 100-subject *Discovery* dataset and then replicated within an 100-subject *Replication* cohort.

Principal component analysis (PCA) was applied to the multi-task BOLD time series, to reorganize the regional BOLD data into a smaller set of spatially orthogonal principal components (PCs; Figure 1A). Across divergent cognitive tasks, we found a dominant, low-dimensional neural signal^10^: the first five PCs accounted for 67.9% of the variance and resolved greater than 90% of the embedding space^17^ (Figure 1B).

**Figure 1:**
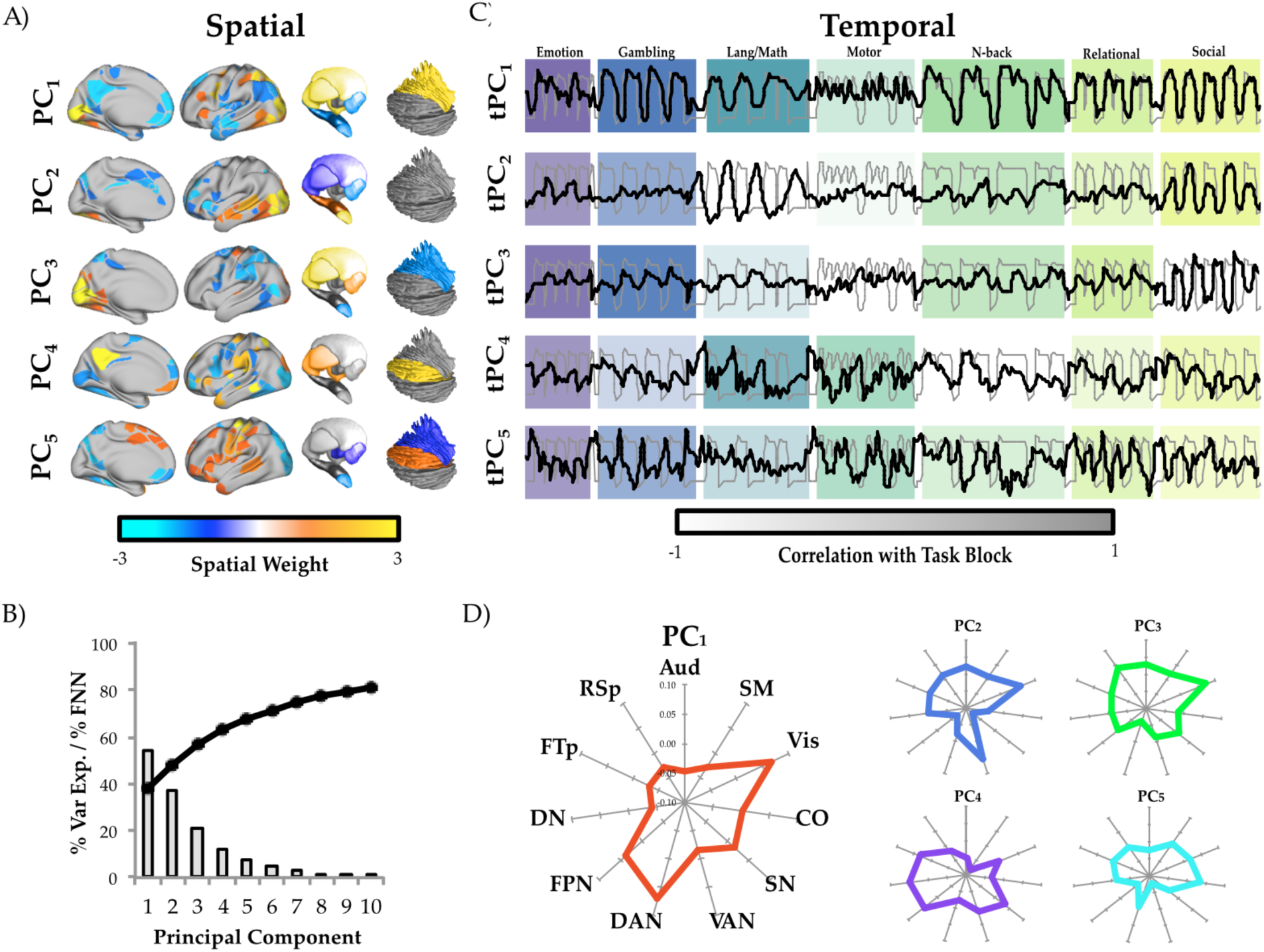
Spatiotemporal principal component analysis across multiple cognitive tasks. A) spatial maps for the first five principal components (colored according to spatial weight; thresholded for visualization); B) line plot representing the percentage of variance explained by first 10 principal components; bar plot depicting the percentage of false nearest neighbors for first 10 principal components; C) correspondence between convolved, concatenated task block regressor (grey) and the time course of the first five tPCs (black): color intensity of the blocks reflect the correlation between tPC_1: 5_ and each of the unique task blocks; D) mean spatial loading of first 5 PCs, organized according to a set of pre-defined networks. Key: DAN: dorsal attention; VIS: visual; FPN: frontoparietal; SN: salience; CO: cingulo-opercular; CPar: cingulo-parietal; VAN: ventral attention; SM: somatomotor; RSp: Retrosplenial; FTP: frontotemporal; DN: default mode; AUD: auditory; tPC: temporal principal component.

The time series of each PC (tPC) was created by weighting the original BOLD time series from the *Replication* dataset with the parcel loading for each spatial component from the *Discovery* dataset at each time point of the experiment. This step allowed us to track the engagement of each PC over time. The first tPC, which explained 38.1% of signal variance across all tasks, reflected a task-dominant signal that was strongly correlated with the overall task block structure across all seven tasks (*r* = 0.64, *p* < 0.01; Figure 1C) and was strongly replicable across cohorts (*r* = 0.84). This result is consistent with previous work that has demonstrated a distributed network of *task-invariant* brain regions that persists across the performance of multiple unique cognitive tasks^4-6^. In addition, the analysis demonstrated prominent spatial overlap between PC1 and dorsal attention, frontoparietal and visual networks, along with striatum, thalamus and lateral cerebellum (Figure 1D). Importantly, fitting a PCA to each of the seven individual tasks separately did not recover the same underlying principal component, but instead identified spatial maps that aligned with the idiosyncratic demands of each task (Figure S1).

The next three components (tPC_2-4_) reflected a closer relationship with specific cognitive tasks (Figure 1C). For example, tPC2 (10.3% variance explained) was associated with the social task and language processing; tPC3 (8.4%) with gambling and emotion; and tPC_4_ (6.3%) with the math task (Figure 1C/D). In contrast, tPC5 (4.8%) was associated with engagement across multiple tasks (*r* = 0.20, *p* < 0.01). The time course of tPC_5_ was correlated with the absolute value of the first derivative of the task design (*r* = 0.11, *p* < 0.01), suggesting that tPC_5_ was uniquely associated with the transition into and out of unique task states. In accordance with this finding, tPC_5_ was associated with activity across a right-lateralized system of cingulate, parietal and opercular cortical regions (Figure 1A) that have previously been shown to play a crucial role in cognitive task engagement and error monitoring^4^.

### Global brain state dynamics

PCA imposes orthogonality onto the data, a crucial step in providing the low dimensional subspace in which to embed the phase space manifold (Figure 2). Other popular methods (such as independent component analysis) find a different set of optimal solutions (such as maximal statistical independence) but these are not, in general, linearly independent^18^. Importantly, state space attractors are invariant to linear transformations of their embedding phase space as long as the dimensions remain orthogonal^19^. Hence, PCA enables analysis of the state-space trajectory (or flow) of the dominant low-dimensional signal, which in turn reflects the temporal evolution of the global brain state. Any residual variance (i.e., from those components not included in the reconstruction) represents a stochastic influence on the ensuing flow.

**Figure 2:**
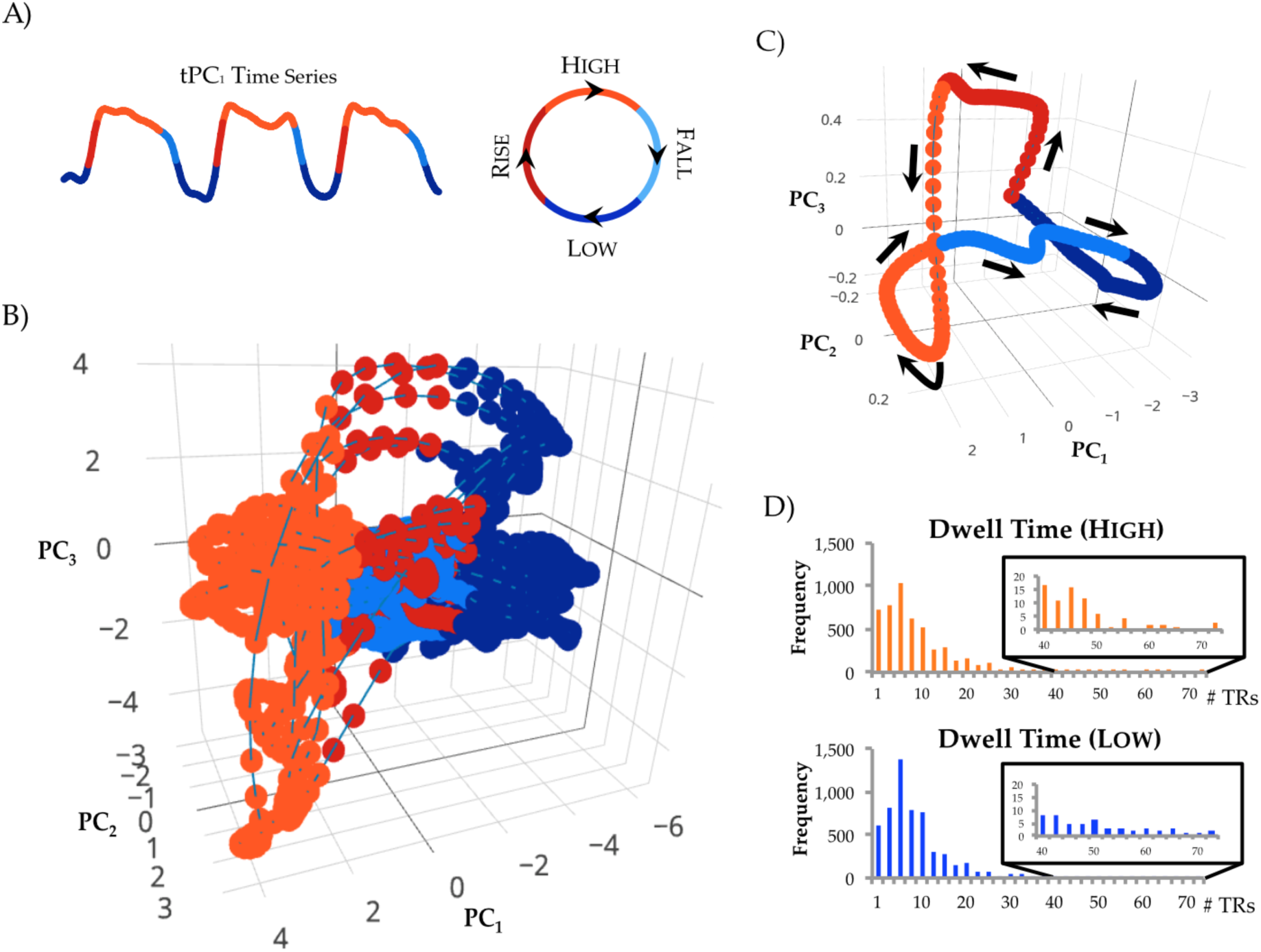
A low-dimensional embedding space characterizes the temporal organization of the brain across cognitive tasks. A) The procedure used to partition tPC_1_ into unique phases – LOW (blue), RISE (red), HIGH (orange) and FALL (light blue); B) a 3-dimensional scatter plot comparing the first three tPCs, colored according to the character of tPC_1_; C) the low dimensional manifold traversed by the global brain state across the first three dimensions, with arrows depicting the direction of flow along the manifold; D) the dwell times for the HIGH and LOW sections of tPC_1_, which were best fit by exponential distributions: inserts demonstrate the tails of each distribution.

To facilitate further analysis of the low-dimensional embedding space, the tPC1 time course was partitioned into relative phase segments^11^: a trough in tPC_1_ defined the LOW phase (blue in Figure 2A); an increase in tPC1 defined the RISE phase (red); a plateau in the tPC_1_ signal defined the HIGH phase (orange); and a decrease in tPC_1_ defined the FALL phase (light blue) of the low-dimensional flow of tPC_1_ (see Methods for details). The resulting phase portrait describes the temporal evolution of the low-dimensional signal shared across all behavioral tasks (Figure 2B and Movie 1; see Figure S2 for projection of tPC_4-5_). The different phases of tPC_1_ were distinctly related to exogenous task demands, with a greater frequency of HIGH phases during task blocks across all seven tasks, and LOW phases during interleaved rest blocks across tasks (*p* < 0.01). By calculating the mean activity across the trajectories in Figure 2B, and projecting these into the embedding space, we were able to recover a canonical low dimensional manifold^20^ that transcends multiple cognitive task states (Figure 2C)^21^ and that was similar following regression of task-mediated effects prior to PCA^22^ (Figure S3). After accounting for task-effects, the distribution of tPC1 dwell times was best described by an exponential distribution (Figure 2D), which is consistent with a noise-driven multistable process^25^, in which the global brain state transitions between relatively shallow attractors. That is, even though the *average* flow is smooth, the *naϊve* (i.e. non-averaged) flow bears the imprint of noise-driven excursions. However, it is important to clarify that the specific nature of the flow through this embedding space invariably reflects a combination of the timing and particular cognitive processes driven by the specific tasks, and thus would not be invariant to changes in external context^23^.

### The cognitive relevance of the global brain state

Having demonstrated that brain state dynamics can be effectively described by the temporal evolution along a low-dimensional trajectory, we were next interested in understanding the potential cognitive relevance of the brain’s functional architecture. Although the seven tasks were qualitatively distinct, we predicted that each of the tasks should recruit similar cognitive capacities, albeit to varying degrees that were defined by idiosyncratic task challenges and complexity.

To test this hypothesis, meta-analytic data was utilized from an existing ‘topic-modeling’ analysis that identified the latent structure present across 5,809 functional neuroimaging studies. This approach links spatial BOLD activation patterns to the ‘topics’ investigated in the original fMRI experiments^26^. Four ‘topic families’ – representing ‘Motor’, ‘Cognitive’, ‘Language’ and ‘Memory’ – were identified by clustering the activation patterns from a 50 topic solution using the NeuroSynth repository (Figure 3A; each family represented a number of sub-topics identified in the meta-analysis; results were confirmed using a ‘reverse inference’ approach; Table S1)^26^. A time series was then created for each topic family by weighting the original BOLD data with the spatial activation pattern of each topic family over time. Comparison with the PC time series revealed a clear relationship between the tPC time series and latent cognitive processes – for instance, ‘Motor’ and ‘Cognitive’ functions were jointly separated from ‘Memory’ and ‘Language’ function by tPC_1_, but were separated from one other by tPC_5_ (Figure 3B). These results confirmed our hypothesis that flow along the low-dimensional manifold was associated with the recruitment of distinct sets of cognitive functions.

**Figure 3:**
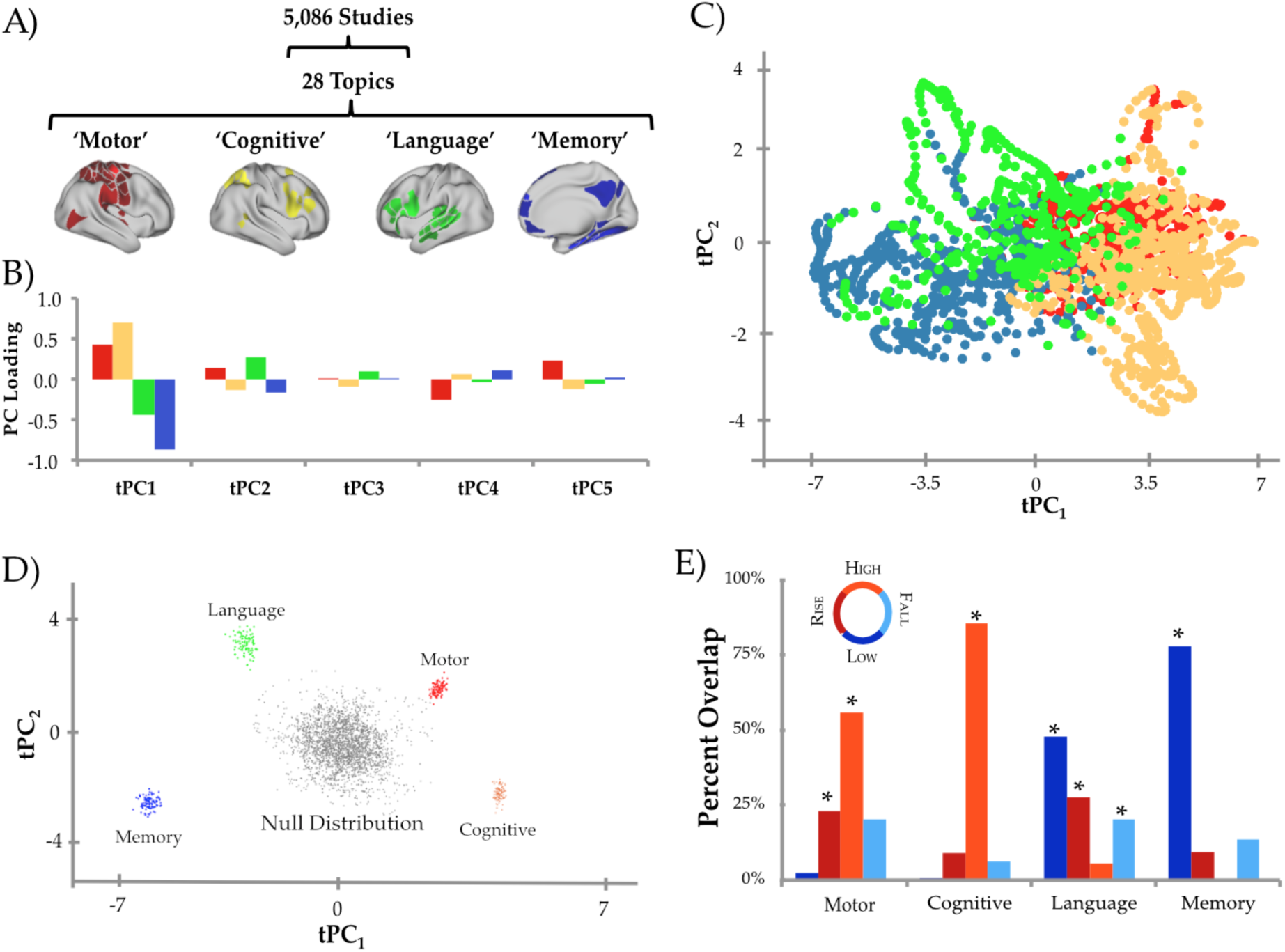
The cognitive relevance of the low-dimensional embedding space. A) Four NeuroSynth “topic families”: Motor (red), Cognition (yellow), Language (green) and Memory (blue); B) bar plot demonstrating loading of Topic Families onto top five principal components; C) scatter plot of time points of the first two tPCs, colored according to their loading onto each of the four NeuroSynth topic families; D) mean value (bootstrapped 100 times) of tPC_1-2_ for each topic family, compared to a block bootstrapped null distribution (5,000 iterations); E) temporal conjunction between the topic families and the four phases of the tPC1 manifold – asterisks denote *p* < 0.01 (block bootstrapped null model).

To project the topic families into the low-dimensional embedding space, the regional BOLD pattern at each time point was assigned to the topic family to which it demonstrated the strongest spatial correspondence using a ‘winner-take-all’ approach (Figure 3C). To test for significance, we constructed a null dataset (5,000 iterations) using a block-randomization bootstrapping procedure that arbitrarily (and repeatedly) splits and reorganizes data over time, similar to the way a dealer would ‘cut’ a deck of cards. This approach scrambles the alignment of the data to the task structure but largely preserves autocorrelation, which can have important influences on the relative degrees of freedom in the data. By comparison to this null distribution, the four topic families occupied unique subspaces of the low dimensional manifold (Figure 3D): the ‘Motor’ and ‘Cognitive’ families were active during HIGH phases, whereas the ‘Memory’ and ‘Language’ families were active during LOW phases of the manifold (*p* < 0.01; Figure 3E). These results highlight the clear relationship between flow on the low-dimensional manifold and recruitment of specific cognitive processes.

### Complex cognitive brain state dynamics

We hypothesized that flow along the tPC_1_ dimension represented an integrative core that balances the competing requirements of global integration (adaptively modifying the functional network signature of the brain in response to task demands) and differentiation (ensuring the distinctive configuration of neural systems required of each cognitive state)^2,27^. To test this hypothesis, we calculated time-varying functional connectivity from the concatenated BOLD time series (after first regressing task effects from each time series) and applied graph theoretical analyses to the resultant temporal connectivity matrices. We used a general linear model to examine the relationship between the tPC_1-5_ time series and time-resolved network architecture, allowing the identification of the topological signature of each PC. Expression of tPC_1_ was associated with a distributed and integrated network topology with strong connections across functionally specialized modules (Figure 4A). In contrast, the topological signatures of lower components were more heterogeneous (Figure 4B). Specifically, tPC_2-4_ demonstrated a trade-off between integration and segregation, whereas tPC_5_ displayed a relatively segregated signature (Figure 4C). These patterns suggest that different low-dimensional components may reflect unique constraints on the balance between integration and segregation in the brain.

**Figure 4:**
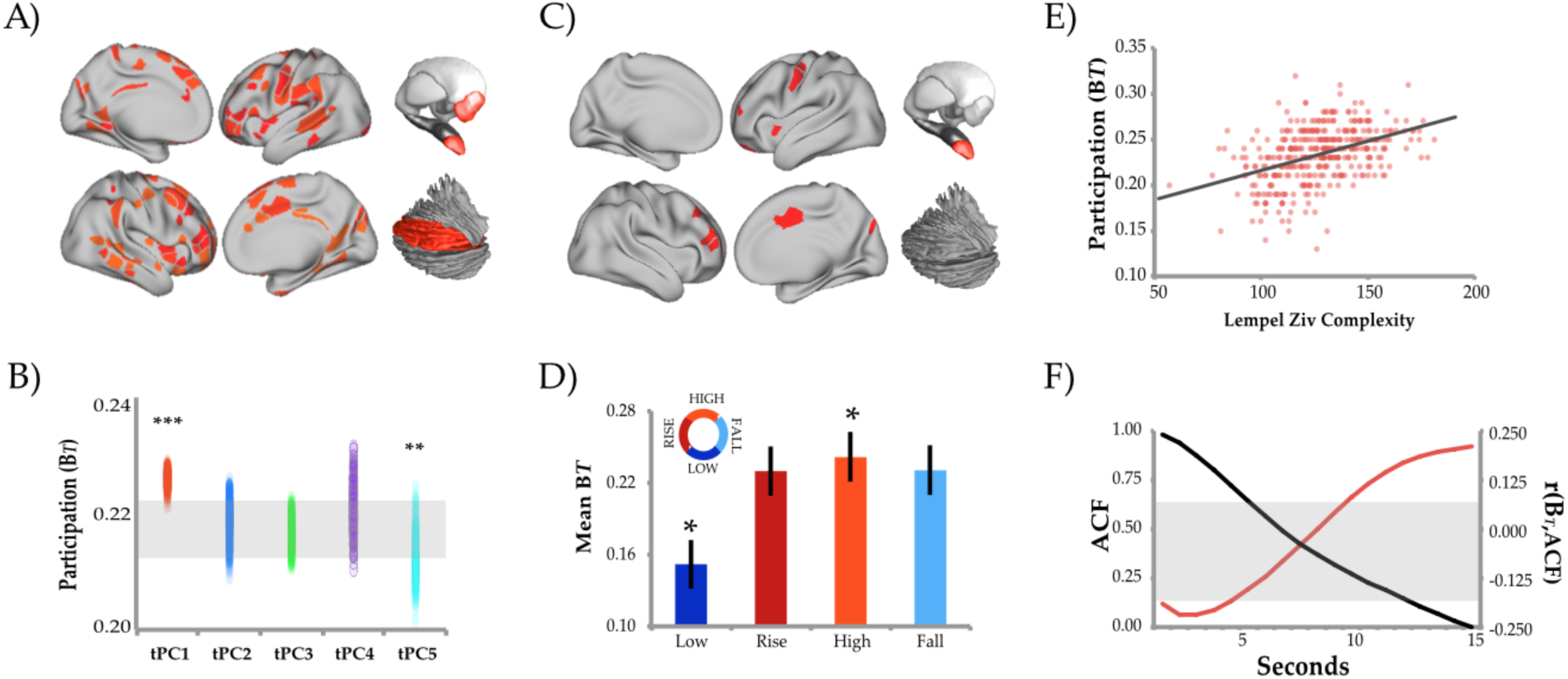
The low-dimensional integrative core of the brain across cognitive tasks. A) surface plot of the most integrative regions in tPC_1_ (regions significantly more integrative than null model); B) parcel-wise participation coefficient (B_*T*_) across the top 5 PCs, with block resampling null values displayed as grey region; stars designate significance of mean B*T* across the whole brain (***-*p* < 0.01; **-*p* < 0.05); C) a surface plot of the most integrative regions in tPC_5_; D) mean B*T* for each of the four phases of the tPC_1_ manifold: asterisks denote *p* < 0.01 (block bootstrapped null model); E) correlation between regional B_*T*_ and Lempel Ziv Complexity (*r* = 0.424; *p* < 0.001); F) black line: mean autocorrelation function (ACF) for all regions; red line: correlation between regional ACF and B_*T*_; grey rectangle – values for which null hypothesis was not rejected.

The most integrative regions associated with PC1 were diffusely distributed across the majority of canonical ‘resting state’ networks, involving regions within the frontal, parietal and temporal cortex, along with the bilateral amygdala and the lateral cerebellum (Figure 4A). There was a distinct relationship between time-resolved network topology and the low-dimensional manifold (Figure 4D): the topological architecture of the brain was more integrated during the HIGH phases (median participation coefficient [B_*T*_] across all parcels = 0.24 ± 0.1; *p* < 0.01) but segregated during the LOW phases (0.15 ± 0.1; *p* < 0.01). There was also a significant relationship between global network integration (mean B_*T*_) and the ‘Cognitive’ topic family (mean B*T* across all parcels = 0.24 ± 0.1; *p* < 0.01). In contrast, the ‘Memory’ topic family was associated with a more segregated network topology (0.15 ± 0.1; *p* < 0.01).

Although system-wide integration is an important signature of complex networks, biological systems also need to retain sufficient flexibility to cope with an array of adaptive challenges. That is, the global brain state should also demonstrate differentiation^2^, reflecting the need for each state of the brain to be distinct from every other possible state. To test this prediction, we estimated the Lempel Ziv complexity of each regional time series^7^. Lempel Ziv complexity estimates the number of distinct binary sequences required to recapitulate a test sequence – information-rich time series have higher complexity and thus require a larger dictionary of sequences in order to recreate them^7^. As predicted^2^, the mean regional signature of integration was positively correlated with Lempel Ziv complexity (*r* = 0.42; *p* < 0.01; Figure 4E).

The brain can also control information flow by modulating inter-regional interactions at different temporal scales. Previous work has demonstrated a heterogeneity of time-scales across the brain^29,30^, in which sensory regions process information quickly (i.e., on the order of milliseconds-to-seconds) whereas more integrated hubs attune to information on slower time-scales (i.e., seconds-to-minutes). To determine whether the low-dimensional topological signature of tPC_1_ was associated with a unique temporal signature, we correlated the extent of autocorrelation within each region with the loading between the time-resolved participation coefficient (B_*T*_) and tPC_1_. This analysis revealed a negative correlation at shorter time scales (0.72 – 5.02 seconds) and a positive correlation at longer time scales (9.36 – 13.68 seconds; *p* < 0.05; Figure 4F), suggesting that tPC_1_ processes information at relatively slow time-scales.

### Neurotransmitter receptor gradients gate cognitive brain dynamics

We next sought to understand the factors that control flow on the low-dimensional manifold outside of the particular sensory constraints imposed by each task. A plausible candidate for orchestrating global control over brain state dynamics are the ascending neuromodulatory systems of the brainstem and forebrain^31^. These highly conserved nuclei project widely throughout the brain to modulate the ‘gain’ of receptive neuronal populations and hence, alter inter-regional communication^32^. That is, they are able to broadly modulate brain network connectivity in a flexible manner^33^. An extensive literature links neuromodulatory systems with to a broad range of cognitive functions^34^, and receptors from several neurotransmitter families have been implicated in either facilitating or inhibiting cognitive processing^31^. Interactions among these systems are also crucial, suggesting that the neuromodulatory system acts as a complex adaptive network that maintains non-linear influence over brain network topology and dynamics^35^.

To test this hypothesis, we used the Allen Brain Micro-Array Atlas (http://human.brain-map.org/) to identify the spatial coverage of a range of metabotropic neurotransmitter receptors using post mortem data on variation in neurotransmitter receptor gene expression. We investigated two main classes of receptor with known opposing effects on cognitive function: a *facilitatory* group, including the dopaminergic D_1_^31^, the noradrenergic α2A^31^, the cholinergic M_1_^36^ and the serotonergic 5HT_2A_ receptors^37^; and an *inhibitory* group, including the D_2_, α1A and 5HT_1A_ receptors (due to inconsistent evidence in the literature, no muscarinic receptors were included in the inhibitory group). Each of these receptors modulates the signal-to-noise ratio in neurons by activating G-protein-coupled receptors^38-40^, with effects typically most pronounced at the network level^41,42^.

To compare brain state dynamics with neuromodulatory coverage, we related the spatial pattern of receptor density to each of the system-wide signatures identified in the preceding analyses (Table S2). We observed opposing relationships between the neuromodulatory groups and the first two tPCs (see Table S2): the facilitatory group loaded positively onto tPC_1_, whereas the inhibitory group loaded negatively (Figure 5). In contrast, tPC_2_ better delineated the receptor families, loading positively onto dopaminergic and noradrenergic, and negatively onto serotonergic and cholinergic receptors (Figure 5). The neuromodulatory groups also demonstrated unique topology: the facilitatory group was associated with increased functional integration (Figure 5; Modularity *[Q]* = 0.14), whereas the inhibitory group was relatively segregated (*Q* = 0.55). Our results thus provide a link between neuromodulatory system heterogeneity and the dynamic neural states required for diverse cognitive tasks^35,43^.

**Figure 5:**
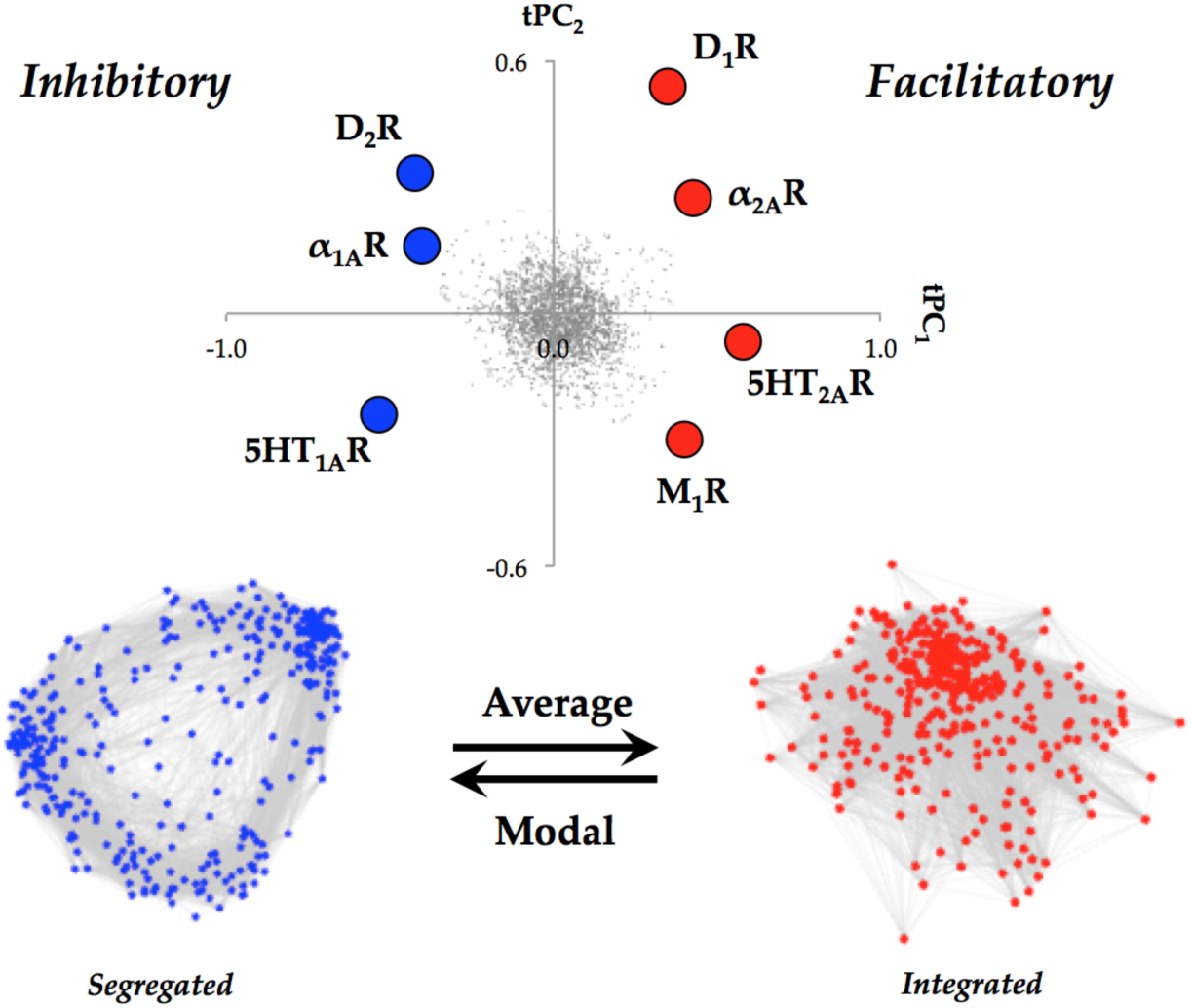
The neurochemical signature of integrated cognitive function. The mean density of tPC_1-2_ for two classes of neurotransmitter receptor maps: a group known to facilitate cognition (D_1_, α_2A_, 5HT_2A_ and M_1_; right) and a group known to inhibit cognition (D_2_, α_1A_, 5HT_1A_; left). The spatiotemporal patterns in these two classes were associated with differential network topologies: the facilitatory group was associated with an integrated brain, whereas the inhibitory group was associated with a relatively segregated brain – force directed plots reflect the mean time-resolved functional connectivity matrix when loading was positive for either the inhibitory (blue) or facilitatory (red) group (thresholded and binarized at 10% density for visualization purposes); and different classes of controllability: the facilitatory states were associated with high average controllability, whereas the inhibitory states were associated with high modal controllability.

The spatial overlap between low-dimensional system dynamics (estimated from task fMRI) and neurotransmitter receptor topography (estimated from *post mortem* brain tissue) suggests that global brain state dynamics may be controlled by the recruitment of distinct neurotransmitter classes^44^. To test this hypothesis, we compared the spatial maps for each receptor sub-type with structural network signatures that mediate distinct control patterns in the human brain: some regions are able to drive the brain into many different states (known as ‘average’ controllability), whereas others facilitate the engagement of ‘hard to reach’ states (‘modal’ controllability)^45^ (Figure 5). The regional signature of these two control classes were estimated using structural diffusion data from the Human Connectome Project^46^, and then related to the receptor maps from the Allen Brain Atlas. We observed strong positive correlations between the *facilitatory* group and ‘average’ controllability, and between the *inhibitory* group and ‘modal’ controllability (Figure 5; Table S2)^45^. Further analysis showed that the spatial loadings for PC^1^ were selectively correlated to each region’s strength (weighted degree; *r* = 0.34; *p* < 0.001). Our results are therefore consistent with the notion that control over network dynamics is a relatively distributed capacity^47^ that may be mediated by highly conserved neuromodulatory systems that guide the flow of the brain across the low-dimensional manifold.

## Discussion

The neural activity required for the execution of cognitive tasks corresponds to flow within a low-dimensional state space. Across multiple, diverse cognitive tasks, the dynamics of large-scale brain activity engage an integrative core of brain regions that maximizes information-processing complexity and facilitates cognitive performance; only to then dissipate as the tasks conclude, flowing towards a more segregated architecture. Our findings further suggest that the brain’s dynamic trajectory is guided by a diverse set of highly conserved modulatory neurotransmitter systems that transition between distinct phases of the attractor, thus providing a plausible biological mechanism for the control of brain state dynamics. Overall, our results provide a novel framework for studying cognitive neuroscience from the perspective of large-scale dynamical systems, linking the flow of cognition to the dynamic reconfiguration of functional networks in the human brain driven by distinct neuromodulatory systems.

The low-dimensional, integrated component that occurs on the plateau phase of the state-space trajectory recurs across multiple unique tasks and forms the functional backbone of cognition within the human brain. Fluctuations in this core network architecture are associated with maximal information processing complexity across relatively long time-scales, suggesting that the temporal signature of the integrative core is information-rich and accumulates information over long time-scales^29,30^. This slow, integrative core – consistent with prior multi-task analyses^4-6^ – contrasts with the architecture during epochs in which there was no task, in which the brain occupied a segregated topology and was associated with a shorter time-scale of information processing. We also find a low-dimensional component corresponding to task onset and offset (tPC_5_). Notably these results were robust to permutation testing and replicated in an independent cohort. However, the specific form of the flow we describe likely reflects idiosyncrasies of the task fMRI battery and in particular, the predominant use of visual stimuli and reliance on motor responses. Nonetheless, we propose that a flow between the integrated and segregated phases will persist beyond the specific tasks used here, and likely constrains cognitive capacities across a variety of psychological contexts. Future experiments could usefully examine the low-dimensional architecture of the brain across a broader range of psychological capacities, including ecologically valid contexts^48^ such as naturalistic stimulation paradigms.

It has been known for some time that ascending neuromodulatory systems provide important constraints on cognitive function^31,49^, but the systems-level mechanisms responsible for these capacities have remained relatively obscure. Here, we provide a mechanistic account of the association between these distinct neuromodulatory systems and cognitive function. Specifically, a diverse set of modulatory neurotransmitter receptors occupy a privileged spatial location in the cortex that maximizes their ability to modulate the flow of cognitive brain states over time^23^. Neuromodulatory receptors stimulate G-protein coupled receptors, which alter trans-membrane ion gradients, and thus make neurons more (or less) likely to fire in response to glutamatergic input^39^. Mechanistically, this process has been interpreted as altering the signal-to-noise ratio within neural circuits^50^ – that is, neuromodulatory receptors play an information-gating role in the brain^51^. Crucially, computational work has also linked neuromodulatory activity to the alteration of the current attractor state^52^, which in turn could influence the flow of low-dimensional activity in the brain to facilitate cognition (Figures 3 and 4). Our results provide empirical evidence for these concepts, and further support the notion that neuromodulation exerts network-level effects on the brain^42,50^. Indeed, it is these non-linear, competitive and cooperative dynamic interactions between neuromodulatory systems that likely imbue the nervous system with its remarkable flexibility^35,43^, enabling the hard-wired “backbone” of the brain to dynamically facilitate the neural coalitions required to navigate an evolving affordance landscape^53^.

Our observations yield novel predictions that can guide future work. For example, similar low-dimensional analyses of brain function in other species, notably other primates, might clarify whether or not shifts in this organizational framework underpin some of the distinctive cognitive abilities of humans. Strong phylogenetic conservatism in neuromodulatory systems suggests that evolution has modified pre-existing structures to shape human cognitive function; and the neural architecture that we have described offers a plausible substrate for such changes. Within our own species, variation in the low-dimensional core of the brain may also underlie some psychological manifestations of neuropsychiatric and neurodegenerative disorders. If so, detailed mapping of individual differences in the integrative core may suggest novel therapeutic interventions. Future work could use the form of the attractor to enable model-based hemodynamic deconvolution, hence uncovering the form of the generative processes. More generally, we hope that this work will provide a platform for future insights into the modular and integrative processes that form the infrastructure for cognition in the human brain.

## Acknowledgments

Data were provided by the Human Connectome Project (HCP), and the Washington University, University of Minnesota, and Oxford University Consortium (Principal Investigators David Van Essen and Kamil Ugurbil; Grant 1U54MH091657) funded by 16 NIH Institutes and Centers that support the NIH Blueprint for Neuroscience Research, and the McDonnell Center for Systems Neuroscience at Washington University. This project also made use of Connectome DB and Connectome Workbench, developed under the auspices of the HCP (http://www.humanconnectome.org/). Neurotransmitter receptor data from the Allen Human Brain Atlas (© 2010 Allen Institute for Brain Science. Available from: human.brain-map.org) were obtained from neurosynth.org. We would like to thank Tim Verstynen for the diffusion data, and Danielle Bassett for the controllability code. The funding for the study was provided by an NHMRC CJ Martin Fellowship (GNT1072403). The study was designed by all authors and conducted primarily by JMS, who also wrote the first draft. All authors provided critical feedback on the manuscript, including editing of the final manuscript.

## Methods

### Data acquisition

Data used in the preparation of this work were obtained from the Human Connectome Project (HCP) database. For both the Discovery and Replication analyses, minimally preprocessed fMRI data were acquired from 100 unrelated participants (mean age 29.5 years, 55% female^54^). For each participant, BOLD data from the LR encoding session from seven unique tasks were acquired using multiband gradient-echo EPI, amounting to 23 minutes 17 seconds of data (1940 individual time points) per subject. The following parameters were used for data acquisition: TR = 720 ms, echo time = 33.1 ms, multiband factor = 8, flip angle = 52 degrees, field of view = 208×180 mm (matrix = 104 × 90), 2×2×2 isotropic voxels with 72 slices, alternated LR/RL phase encoding.

### Data pre-processing

Bias field correction and motion correction (12 linear DOF using FSL’s FLIRT) were applied to the HCP resting state data as part of the minimal preprocessing pipeline^54^. Temporal artifacts were identified in each dataset by calculating framewise displacement from the derivatives of the six rigid-body realignment parameters estimated during standard volume realignment^55^, as well as the root mean square change in BOLD signal from volume to volume (DVARS). Abnormal frames were not excluded from the data. However, we observed no significant relationship between any of the tPC time series and framewise displacement (estimated from the temporal head motion parameters) at the individual subject level (*p* > 0.5). Following artifact detection, nuisance covariates associated with the 12 linear head movement parameters (and their temporal derivatives), frame-wise displacement, DVARS, and anatomical masks from the CSF and deep cerebral WM were regressed from the data using the CompCor strategy^56^. To ensure equivalence across tasks, the data were also normalized within each temporal window, which effectively controlled for the global signal, while also equilibrating the data across independent subjects. Finally, a temporal low-pass filter (*f* < 0.125 Hz) was applied to the data.

### Brain parcellation

Following pre-processing, the mean time series was extracted from 375 pre-defined regions-of-interest (ROI). To ensure whole-brain coverage, we extracted 333 cortical parcels (161 and 162 regions from the left and right hemispheres, respectively) using the Gordon atlas^16^, 14 subcortical regions from Harvard-Oxford subcortical atlas (bilateral thalamus, caudate, putamen, ventral striatum, globus pallidus, amygdala and hippocampus), and 28 cerebellar regions from the SUIT atlas^57^.

### Cognitive tasks

Seven unique tasks were utilized from the Human Connectome Project consortium: Emotion, Gambling, Language/Math, Motor, N-back, Relational and Social^15^. For each task, a block regressor was created by partitioning the time series into time points in which subjects were actively performing the task; and those in which they were ‘resting’ (note: not all tasks contained designated ‘rest’ blocks). Of note, the ‘rest’ blocks in the Math task involved an auditory, language-based task. The time points associated with each block were convolved with a canonical haemodynamic response function (using the *spm_hrf.m* from SPM12) and then concatenated over time to create a single task block regressor. These served as reference time series for comparison to the tPC time series (see grey lines in Figure 1B). All results were successfully replicated using a finite impulse response model.

### Principal component analysis

Pre-processed data from each task were concatenated to form a multi-task time series per subject and a spatial PCA was performed on the resultant data^58^. Task block structure was not regressed from the data prior to PCA estimation. The time series of each PC was then estimated by calculating the weighted mean of the group-level BOLD time series associated with each respective PC^11^. To aid inference, group-level tPC time series were calculated by taking the mean for each PC time series across all subjects. To estimate the appropriate dimensionality of the data^17^, we calculated the percentage of false nearest neighbors following the PCA decomposition (a measure of effective embedding across dimensions^17^), and found that there were < 10% present in the top five PCs, and <1% present in the top ten PCs (Figure 1B).

To ensure that the low-dimensional embedding space was not adversely affected by the task block structure, we replicated our analysis using the residuals of a ordinary least squares regression in which we modeled regional BOLD data according to the task structure present across all seven tasks (with independent blocks modeled as unique regressors). Each of the major results in our study was replicated following this procedure, suggesting that, although the low-dimensional signature of the brain was related to the temporal structure imposed by the tasks, this factor was not solely responsible for the psychophysiological relationships observed in our study.

To ensure that the PCA results were robust to individual differences, we ran two subsequent analyses: i) the same analysis was conducted in a Replication cohort (N = 100); and ii) a bootstrap analysis was performed by estimating a PCA on 100 randomly chosen subjects (with replacement, from a larger pool of 200 unrelated subjects). Each of these analyses was associated with robustly similar results to the group mean analysis (*r* > 0.85).

To determine the importance of running the PCA across all concatenated tasks, we performed three subsequent analyses: i) we re-ran separate PCAs for each task, individually and found that, although one of the first five components for each task was strongly related to task block structure (*r* > 0.50), the spatial weightings were dissimilar to the pattern observed when all tasks were concatenated (mean *r* = 0.18 ± 0.2) and more similar to the main effect of each task (mean *r* = 0.72 ± 0.2). A difference score was then calculated between the two groups, which allowed us to estimate statistical significance using the Dunn and Clark Statistic (*ZI\**); ii) we used a bootstrapping approach in which we randomly selected between 2-6 tasks and re-ran the PCA, and then performed both a spatial and temporal correlation between the topography and time series for PC1, respectively. In doing so, we found that at least 4 tasks were required to recreate the pattern found across the original 7 tasks (Figure S1); and iii) we ran a standard GLM using the concatenated task block structure, demonstrating a selective positive correlation between the spatial map of PC_1_ and the resultant Beta map (*r* = 0.94; *p* < 0.001). A similar significant relationship was selectively observed between PC_5_ and the Beta map estimated from a GLM comparing task-onsets to BOLD activity (*r* = 0.34; *p* < 0.001).

### Relationship between principal components and task block structure

To estimate the relationship between the tPC time series and the task block structure, we ran a series of linear regression analyses comparing the temporal fluctuations in tPC time courses and the concatenated, convolved task block time series, both for the entire set of seven tasks and also for each task independently. In a similar fashion to the spatial PCs, tPCs were strongly replicable across the Discovery and Replication cohorts (mean *r* across tPC_1-5_: 0.84 ± 0.1), confirming the specificity of low dimensional temporal brain activity during cognitive task performance. Finally, none of the tPCs were significantly correlated with typical noise confounds, such as head motion (framewise displacement; mean *r* across tPC_1-5_: 0.00 ± 0.1) or signals from the white matter and cerebrospinal fluid (mean *r* across tPC_1-5_: 0.00 ± 0.1) at the individual subject level, nor the global signal over time (*r* = −0.04). In addition, there were no interactions between task blocks and head motion, and results were found to be replicable when performing moderate levels of ‘scrubbing’ (i.e., censoring data with framewise displacement > 0.25 and DVARS > 2.5%)^2^.

### Low-dimensional manifold

To describe flow along the low-dimensional embedding space, we used an approach previously utilized in *C. elegans* calcium imaging^11^, in which the global brain cycle is partitioned into four ‘phases’ using the time course of the tPC_1_ as a reference signal: a trough in tPC1 defined the LOW phase (tPC_1_(t) < 33^rd^ percentile; blue in Figure 2A), an increase in tPC1 defined the RISE phase (33^rd^ percentile < tPC_1_(t) < 67^th^percentile & dtPC_1_’ > 0; red), a plateau in the tPC_1_ signal defined the HIGH phase (tPC_1_(t) > 67^th^ percentile; yellow), and a decrease in tPC_1_ defined the FALL phase (33^rd^ percentile < tPC_1_(t) < 67^th^ percentile & dtPC_1_’ < 0; green). After classifying low-dimensional activity into these four phases, we then performed a linear interpolation on each trajectory (i.e., to warp each segment into a set of identically-sized vectors). We were then able to estimate the trajectory of a low-dimensional ‘manifold’^20^ by calculating the mean activity across the interpolated trajectories, which in turn could be projected into the embedding space to describe the manifold (Figure 2C).

Using the four tPC_1_ phases (LOW, RISE, HIGH and FALL), we estimated the ‘dwell time’ for each of the phases explored by the first PC by calculating the number of consecutive TRs in which each phase was present in the data. We then separately fit exponential, Weibull (stretched exponential), power law and gamma distributions to these data (these reflect the likelihood of a multistable, critical or metastable process respectively, see ^25^). The log likelihood of each fit was then used to compute the Bayesian Information Criterion (BIC) for each distribution – low values here represent stronger evidence for a particular fit. We found that exponential fits were better able to explain the data (i.e. lower BIC = −34,637) than were the Weibull (BIC = −33,814), gamma (BIC = −33,620) or power-law (BIC = −7,012) distributions.

### Topic mapping

To determine the potential cognitive relevance of the low-dimensional embedding space, we created regional estimates of 28 spatial maps that represented a curated selection of the 50 ‘topics’ identified during a large-scale analysis of existing neuroimaging literature^26^, in which we excluded topics that were not explicitly related to psychological constructs. These 28 maps were further collapsed into four tight-knit ‘topic families’ (see Table S1) by calculating the spatial similarity of each map and then clustering the matrix using a weighted version of the Louvain algorithm (i.e. the algorithm described in *Time-resolved network topology*). Topic Families were assigned labels according to the top 10 terms associated with the topic-word loading matrix that related study terms to brain mappings. We then created a weighted mean between each of these topic family spatial maps and the concatenated BOLD time series data. Using a ‘winner-take-all’ approach, we categorized each time point according to the topic family with the strongest spatial correspondence to the regional BOLD pattern present at that time, which allowed us to then project the topic families into the low dimensional embedding space (Figure 3C). Finally, we used a non-linear, block bootstrapping permutation test, which preserves some of the autocorrelation structure, to demonstrate that the four topic families were selectively associated with unique aspects of the low dimensional manifold (5,000 iterations; *p* < 0.01).

### Time-resolved functional connectivity

To estimate functional connectivity between the 375 ROIs, we used the Multiplication of Temporal Derivatives (*M*) technique^59^. *M* is computed by calculating the point-wise product of temporal derivative of pairwise time series (Equation 1). The resultant score is then averaged over a temporal window, *w,* in order to reduce the contamination of high-frequency noise in the time-resolved connectivity data. A window length of 20 TRs was used in this study, though results were consistent across a range of *w* values (10-50 TRs). To ensure relatively smooth transitions between each task, connectivity analyses were performed on each individual task separately, and were subsequently concatenated. In addition, all analyses involving connectivity (or the resultant topological estimates) incorporated the junction between each task as a nuisance regressor.

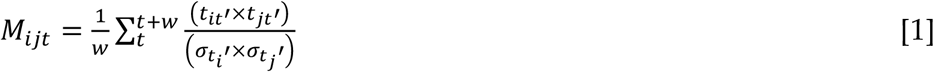

Where for each time point, *t*, the *M* for the pairwise interaction between region *i* and *j* is defined according to equation 1, where *t’* is the first temporal derivative (*t+1* – *t*) of the *i*^th^ or *j*^th^ time series at time *t*, σ is the standard deviation of the temporal derivative time series for region *I* or *j* and *w* is the window length of the simple moving average. This equation can then be calculated over the course of a time series to obtain an estimate of time-resolved connectivity between pairs of regions.

### Time-resolved network topology

The Louvain modularity algorithm from the Brain Connectivity Toolbox (BCT^60^) was used in combination with the MTD to estimate time-resolved community structure. The Louvain algorithm iteratively maximizes the modularity statistic, *Q*, for different community assignments until the maximum possible score of *Q* has been obtained (see Equation 2). The modularity estimate for a given network is therefore a quantification of the extent to which the network may be subdivided into communities with stronger within-module than between-module connections.

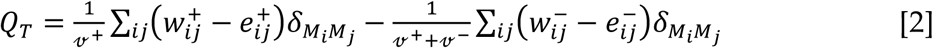

where *v* is the total weight of the network (sum of all negative and positive connections), *w*_*ij*_ is the weighted and signed connection between regions *i* and *j, e_ij_* is the strength of a connection divided by the total weight of the network, and *δ*_*MiMj*_ is set to 1 when regions are in the same community and 0 otherwise. ‘+’ and ‘–’ superscripts denote all positive and negative connections, respectively.

For each temporal window, we assessed the community assignment for each region 500 times and a consensus partition was identified using a fine-tuning algorithm from the Brain Connectivity Toolbox (BCT, http://www.brain-connectivity-toolbox.net/). This afforded an estimate of both the time-resolved modularity (*Q*_*T*_) and cluster assignment (*CiT*) within each temporal window for each participant in the study. We calculated all graph theoretical measures on un-thresholded, weighted and signed connectivity matrices^60^ and the γ parameter was set to 1. Consistent with previous studies^61^, the average number of communities identified in each window was 2.74 ± 0.5.

Based on time-resolved community assignments, we estimated within-module connectivity by calculating the time-resolved module-degree Z-score (W*_T_;* within module strength) for each region in our analysis (Equation 3)^62^, where κ*_iT_* is the strength of the connections of region *i* to other regions in its module s*_i_* at time *T*, 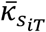 is the average of κ over all the regions in s*_i_* at time *T*, and 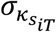 is the standard deviation of κ in s*_i_* at time *T*.

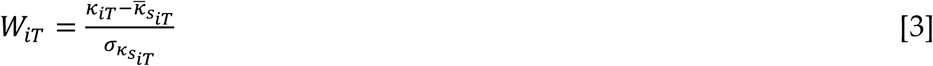

The participation coefficient, *B*_*T*_, quantifies the extent to which a region connects across all modules (i.e. between-module strength) and has previously been used to successfully characterize hubs within brain networks (e.g. see ^63^). The B*T* for each region was calculated within each temporal window using Equation 4, where κ*_isT_* is the strength of the positive connections of region *i* to regions in module *s* at time *T*, and κ*_iT_* is the sum of strengths of all positive connections of region *i* at time *T*. Negative connections were discarded prior to calculation. The participation coefficient of a region is therefore close to 1 if its connections are uniformly distributed among all the modules and 0 if all of its links are within its own module.

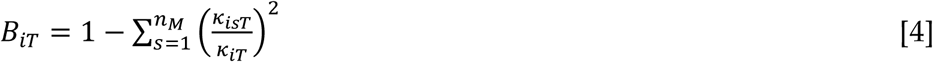

To determine the topological signature of each tPC, we used a general linear model to fit the top five tPC time series to time-varying network topology. Although synchronous sensory inputs do not necessarily force regions to couple together over time^5^, we first regressed out all unique patterns associated with the task blocks from each of the seven tasks (with each unique block modeled as a separate regressor). In addition, the calculation of the temporal derivative also down-weights the task effects on overall activity that are presumed to drive spurious functional connectivity^59^. We then fit a general linear model using the tPC time series as predictors, along with separate regressors that modeled the junction between corresponding tasks. Separate models were fit for unique regional B*_T_* values. We subsequently ran a block bootstrapping permutation test to determine whether there were significantly elevated values of B*T* within predefined networks of the brain (5,000 iterations; *p* < 0.01). In addition, we also analyzed multiple separate window lengths (10 – 100 in steps of 10), and observed a similar relationship between mean B*_T_* and tPC*_1_*, albeit with a peak in similarity at a window length of 20 TRs).

### Complex cognitive brain state dynamics

To explore the functional signature of the integrative core, we calculated the mean intra-regional connectivity for all regions connected to the integrative core (mean regional-to-core connectivity [transformed using Fisher’s r-to-Z] > 1.0) and compared this value to the mean connectivity for all regions outside of the core (i.e. mean regional-to-core connectivity ≤ 1.0). Prior to calculating the difference score, we first applied a Fisher’s r-to-Z transform to each data point to increase Gaussianity. These values were compared using an independent samples t-test.

To estimate time series differentiation, we calculated the Lempel Ziv complexity^7^ of each region’s concatenated time series, binarized to values greater than or less than 0. We then ran a Pearson’s correlation comparing the LZ complexity scores with the participation coefficient (B*_T_*) associated with tPC*_1_* (i.e. the regional beta weights from a general linear model in which the tPC time series were regressed against time-resolved B*T* values). The autocorrelation function was estimated for each region by calculating the time-delayed Pearson’s correlation between each region’s pre-preprocessed BOLD time series, using a lag of 1-30 TRs (0.7 – 21.6s). For each lag, a Pearson’s correlation was conducted between the integrative core and the autocorrelation function of each region. For each analysis, a block resampling permutation test was conducted to test statistical significance.

### Neurotransmitter receptor mapping

To investigate the potential pharmacological correlates of progressive evolution along the manifold, we interrogated the neurotransmitter receptor signature of each region of the brain. To do so, we used the Allen Brain Atlas micro-array atlas (http://human.brain-map.org/) to identify the regional signature of genetic expression of metabotropic neurotransmitter receptors that were *a priori* related to cognitive function. We identified neurotransmitter receptor maps for receptors from four major neurotransmitter families, which were grouped into two families: a facilitatory group, comprising dopaminergic (D*_1_*), noradrenergic (α_*2A*_), cholinergic (M*_1_*) and serotonergic (5HT_2A_) receptors; and an inhibitory group, comprising D2, α1A, and 5HT_1A_ receptors. We first identified the spatial topography of each receptor sub-type. We then created a weighted mean between each of these neurotransmitter receptor maps and the concatenated BOLD time series data. These time series were then related to: a) the tPC time series; b) the topic map time series; and c) the time-resolved topological time series. We applied a series of block-resampling permutation tests to test for temporal alignments between neurotransmitter receptor maps and the manifold and topic family maps, separately (5,000 iterations; *p* < 0.01).

### Structural Controllability

A structural connectome was created from diffusion MRI data from 842 subjects (372 males and 470 females, age 22 ∽ 36) from the HCP cohort using a deterministic fiber tracking algorithm that leverages information in spin distribution functions (for details, see ^46^). The spatial resolution was 1.25 mm isotropic, TR was 5500 ms, TE was 89.50 ms, the b-values were 1000, 2000, and 3000 s/mm2, and the total number of diffusion sampling directions was 90, 90, and 90 for each of the shells, in addition to 6 b0 images. A weighted connectivity matrix was quantified using the same cortical and subcortical parcellation used in the functional analysis. The strength (i.e. weighted degree) of each region was collected for further analysis, and a simple randomized null model (5,000 permutations) was run in order to determine whether the core regions demonstrated greater structural inter-connectivity than the rest of the brain.

To estimate regional controllability, we calculated the average and modal controllability of the weighted structural connectome (see ^45^ for details). Briefly, average controllability is defined as the 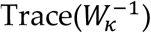, where 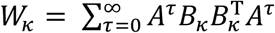 is the Controllability Gramian, *A* is the weighted connectivity matrix and *B* is the input matrix that defines the control points in the network; and modal controllability is computed as the eigenvector matrix *V* = [v_ij_] of the network adjacency matrix *A* (if the entry *v*_ij_ is small, then the *j*^*th*^ mode is poorly controllable from node *i*. While it is known that these measures relate to degree/strength, there is also evidence that they remain after controlling for degree, and hence may relate to other topological features of the structural connectome^47^. The regional patterns created from these analyses were then used to create a weighted mean between each of these control maps and the concatenated BOLD time series data. These time series were then related to the other outcome measures in our study, and we used a block-resampling null model to determine statistical significance. We also correlated the spatial loading of the first five PCs with the strength (i.e. weighted degree) of each region within the structural connectome.

## Supplementary Figures S1-S2

**Figure S1:**
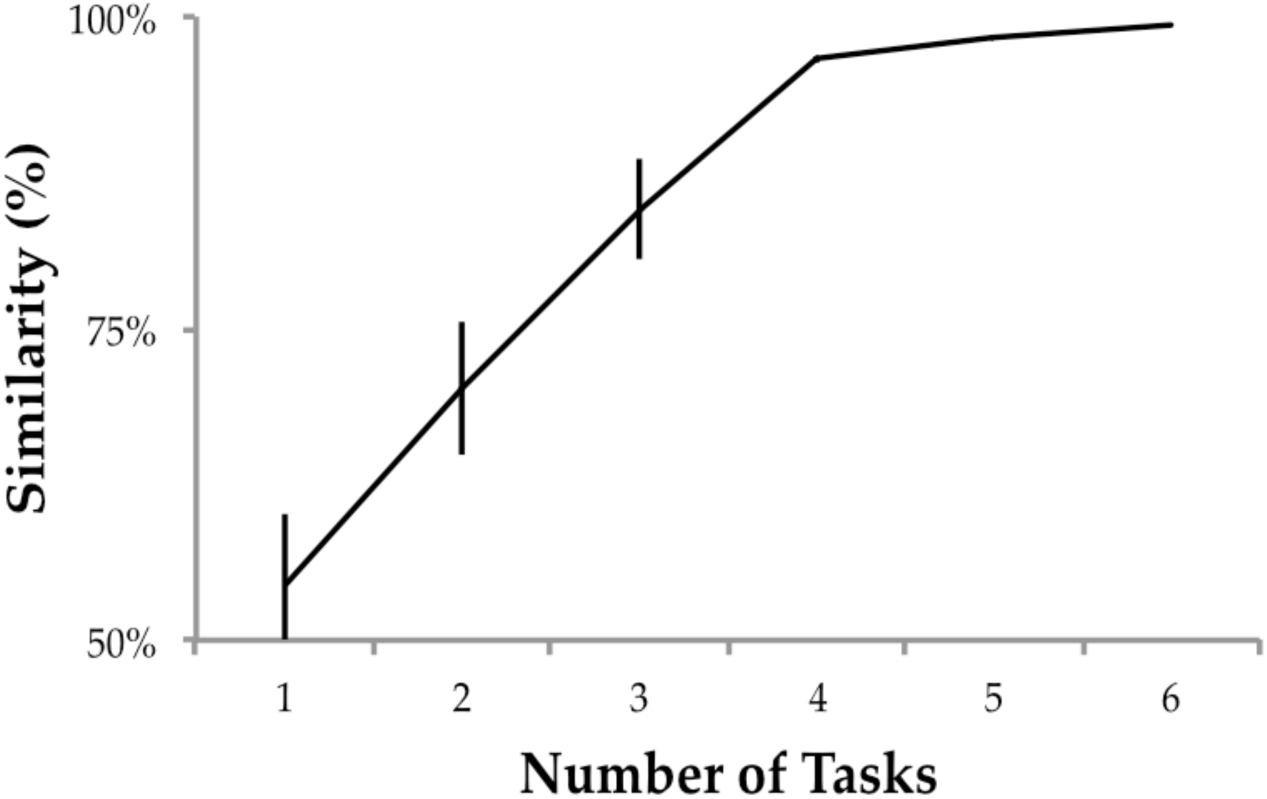
results of the bootstrapping analysis that shows that 4-5 tasks are required to discover the same underlying principal component (tPC_1_) that recurs across task blocks; error bars denote standard error across 100 iterations.

**Figure S2:**
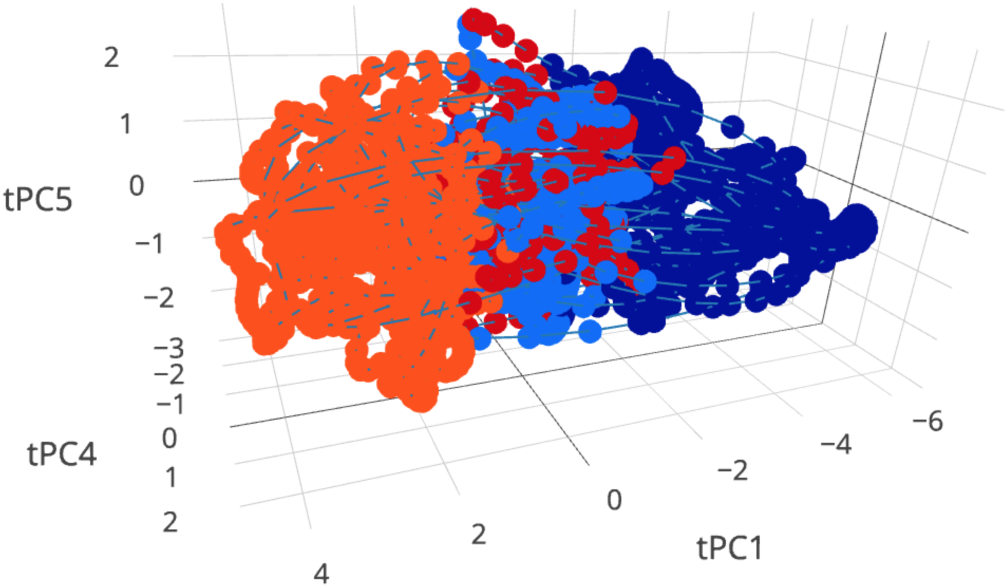
embedding space, comparing tPC_1_ with tPC_4_ and tPC_5_.

**Figure S3:**
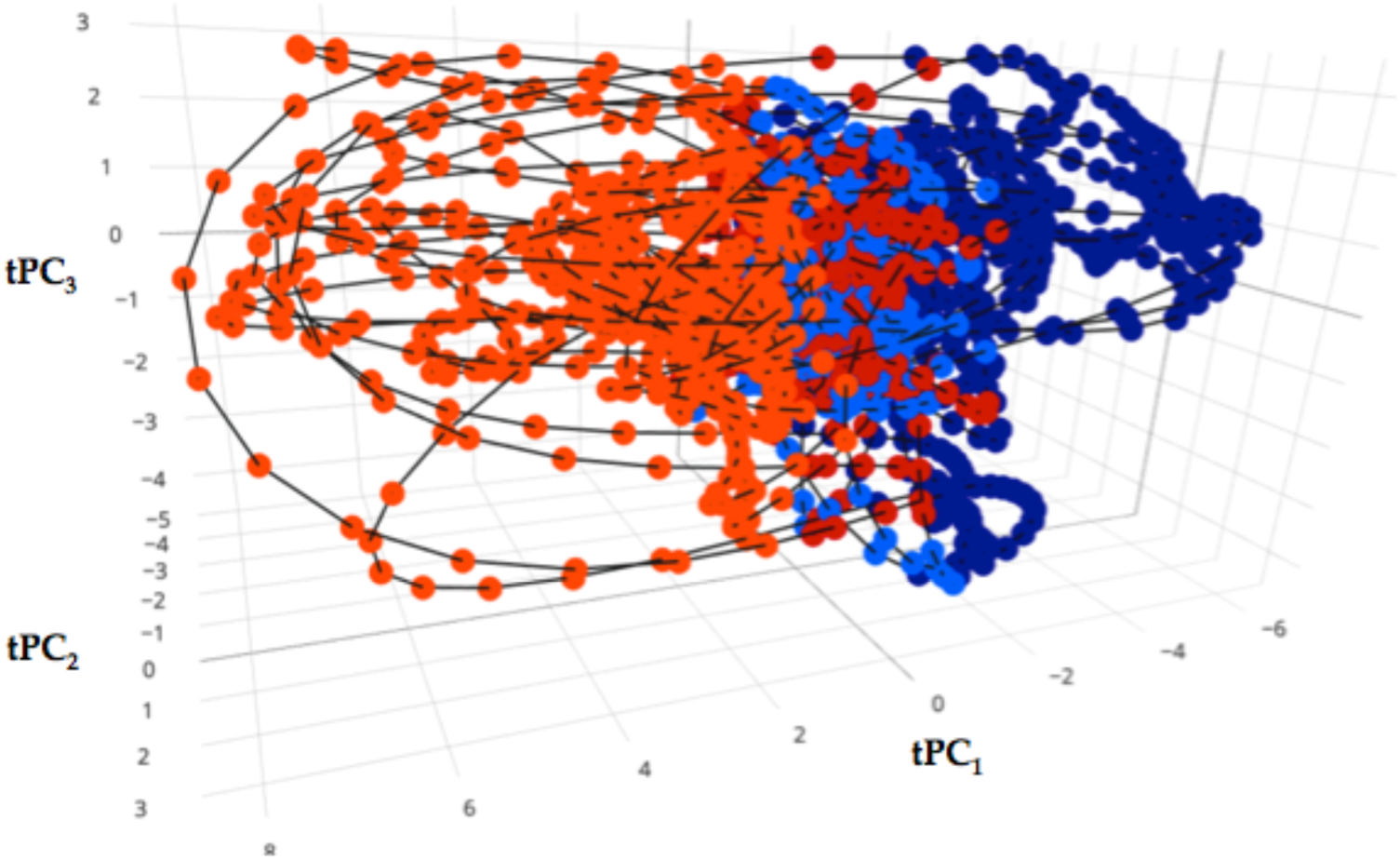
embedding space (tPC_1-3_) following regression of the task block structure.

**Table S1:**
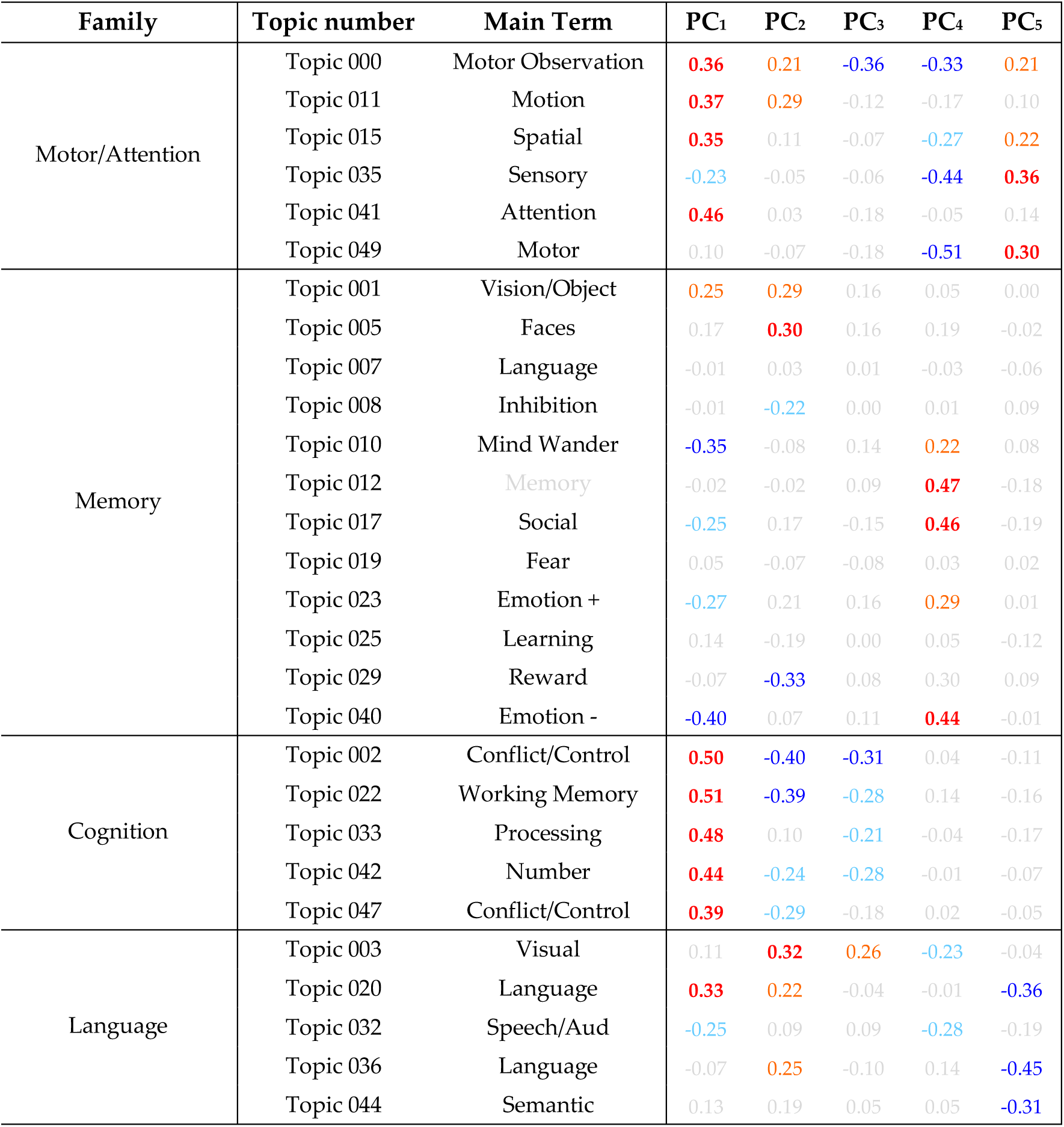
NeuroSynth Topic Families. Values denote Pearson’s correlations: red/blue – p < 0.001; orange/light blue – p < 0.01; grey – p > 0.05.

**Table S2:** Neuromodulatory Families. Values denote Pearson’s correlations: red/blue – *p* < 0.001; orange/light blue – *p* < 0.01; grey – *p* > 0.05.

| Family | Receptor | PC<sub>1</sub> | PC<sub>2</sub> | PC<sub>3</sub> | PC<sub>4</sub> | PC<sub>5</sub> | Mot | Cog | Mem | Lang | B<sub>T</sub> |
| --- | --- | --- | --- | --- | --- | --- | --- | --- | --- | --- | --- |
| Facilitatory | D<sub>1</sub>R | 0.35 | 0.54 | 0.36 | 0.00 | 0.16 | 0.23 | 0.51 | -0.22 | 0.57 | 0.18 |
|  | α<sub>2A</sub>R | 0.43 | 0.27 | -0.24 | -0.28 | 0.22 | 0.26 | 0.39 | -0.30 | -0.10 | 0.19 |
|  | M<sub>1</sub>R | 0.40 | -0.30 | -0.35 | 0.32 | 0.03 | 0.13 | 0.55 | 0.17 | -0.05 | 0.12 |
|  | 5HT<sub>2A</sub>R | 0.58 | -0.07 | 0.23 | 0.04 | 0.12 | 0.75 | 0.80 | -0.35 | 0.31 | 0.18 |
| Inhibitory | D<sub>2</sub>R | -0.42 | 0.33 | -0.06 | -0.05 | -0.05 | -0.51 | -0.71 | 0.32 | 0.07 | -0.17 |
|  | α<sub>1A</sub>R | -0.40 | 0.16 | -0.46 | -0.24 | 0.09 | -0.36 | -0.51 | 0.18 | -0.36 | -0.20 |
|  | 5HT<sub>1A</sub>R | -0.53 | -0.24 | -0.15 | 0.15 | -0.14 | -0.71 | -0.69 | 0.53 | -0.21 | -0.18 |

